# Inter-Tools: a toolkit for interactome research

**DOI:** 10.1101/150706

**Authors:** Hannah Catabia, Caren Smith, José Ordovás

**Affiliations:** JMUSDA Human Nutrition Research Center on Aging, Tufts University, Boston, 02111, USA.; IMDEA Alimentación, Madrid, Spain.; CNIC, Madrid, Spain.

## Abstract

Interactome analysis is an increasingly useful tool for discovering disease pathways. However, protein interaction data is scattered among different sources, and researchers must often sift through several large repositories before beginning a project. Inter-Tools allows users to search databases in the PSI-MITAB standard format for interactions meeting specified criteria, then combine them into a single, species-specific interactome dataset. It also produces graph files of interactome networks and performs basic analysis on user-input gene sets. Inter-Tools reduces the barriers to preliminary interactome investigation and hypothesis generation.

**Availability:** www.github.com/catabia/inter-tools

Supplementary information available at the website listed above.

## 1 Introduction

Interactome analysis is an important method for understanding disease pathways in the genome. However, researchers face many challenges when working with interactome data. The first is creating a customized, up-to-date interactome dataset tailored to the needs of a particular project. There are currently several different repositories that catalog protein-protein interactions (PPIs) in the standard PSI-MITAB [1] format: BioGRID [2], IntAct [3], the Database of Interacting Proteins (DIP) [4], etc. Many contain PPI data for several different species, and it is up to the researcher to extract data for the species being studied. Also, each database contains unique PPI data, as well as data that overlaps with other repositories. Combining several databases yeilds the most complete picture of the interactome possible. Since many PPI databases are regularly updated with new interactions, combining them once is not sufficient. Some databases contain non-experimentally verified interactions; generally, these should be removed to maintain a data quality standard. Finally, different databases use different identifiers for the interacting genes: Entrez [5], UniProt [6], etc. A unified database needs to standardize and convert disparate gene identifiers.

The second challenge to interactome research is visualizing a small, local interaction network within the larger human interactome. There are several graph visualization software programs, such as Gephi [15] and Cytoscape [14], that can be used to inspect interactome data. However, in order to use these programs to their full potential, interactome and disease gene data need to be combined and processed into a format that the software can understand. Inter-Tools streamlines all of the abovementioned tasks with a command-line accessible Python script.

## 2 Tool Description and Functionality

Interact-Tools contains two separate command-line tools written in Python: inter-build.py and inter-map.py. They both require Python 2.7 and R to be installed. A third script, inter-install.py, may be used to automatically download and install all packages and APIs required to run Inter-Tools.

### 2.1 Inter-Build: Assembling an Interactome Databse

Inter-Build converts an unlimited number of MITAB files into a single, unified interactome database for use with Inter-Map. MITABs use a diverse set of gene identifiers. Inter-Build extracts the gene identifiers for each interaction, then uses MyGene API [9] to translate them into Entrez ID [5], UniProt [6], and NCBI Symbol for the final dataset. Users may input one or multiple NCBI Taxonomy IDs to create an interactome for a specific species (default: 9606 - *Homo sapiens).* [10] They may also restrict their dataset to a specific class of interaction detection methods from PSI-MI ontology (default: 0045 - experimental interaction detection). [1] Inter-Build uses OntoCAT API [11] via R to access the PSI-MI interaction detection ontology [1].

Inter-Build creates four files: (a) a CSV file that contains the final interactome dataset, (b) a text file that contains summary information about the dataset, such as the number of genes and interactions, the diameter of the network, etc., (c) a Venn diagram representing the overlap of the three largest MITAB files used to create the interactome, and (d) a histogram that plots the number of interactions per gene.

### 2.2 Inter-Map: Visualizing Gene Sets on the Interactome

Inter-Map creates graph files of the interaction network between a user-input set of genes. The program treats the gene set as a subset of nodes within the interactome. Inter-Map is set up to run seamlessly with an interactome dataset produced with Inter-Build; however, with some simple file modifications described in the Inter-Tools user guide (supplementary materials), it can also utilize PPI datasets from other sources, such as BioPlex 2.0. [12] Inter-Map builds a graph using NetworkX [13] and produces two GEXF files compatible with graph visualization software such as Cytoscape [14] and Gephi [15]. The first is a graph of the full interactome, with the nodes related to the user-input gene set labeled; the second is a graph of *only* the user-input gene set and the interactions between them.

In both GEXF files, each node from the user-input gene set has a weight attribute. This weight describes the ratio of the number of interactions of the gene within the gene set to the number of its interactions in the interactome as a whole. This attribute may be useful for highlighting genes that are specifically well-connected within the geneset, as opposed to generally promiscuous. Inter-Map also produces a RESULTS.txt file that contains information about the genes not found in the interactome, the largest connected component (LCC) of the gene set, and the mean shortest distance between genes in the set (as defined by Menche, *et. al.*). [16]

## 3 Illustrative Example

On April 24, 2017, we downloaded the most recently updated MITAB files from BioGRID [2], IntAct [3], and DIP [4]. We then used Inter-Build to create a human interactome using the default taxid (9606: *Homo sapiens*) and interaction detection method ancestor (0045: experimental interaction detection). The resulting human interactome database has 16,972 unique genes and 273,200 unique interactions. Of these interactions, 147,881 (54.1%) appeared only in BioGRID and 50,692 (18.6%) appeared only in IntAct. Using BioGRID alone would have provided only 80.1% of the human interactome accessible by combining all sources; IntAct alone would have given only 44.7%. The most promiscuous gene in the interactome has 2,091 interactions, while the median number of interactions per gene was 13. The diameter of the entire gene network was 8. (This interactome dataset is available in the supplementary materials, and it is further described in the Inter-Tools user manual.)

We then used this human interacome dataset and Inter-Map to analyze a set of 157 genes identified by the Global Lipids Consortium to be associated with high density lipoprotein cholesterol (HDL), low density lipoprotein cholesterol (LDL), triglycerides, and total cholesterol concentration in humans. [17] A total of 145 (92.4%) of these genes were found in the interactome, 46 of which formed the largest connected component (LCC) of the gene network (Figure 1). The mean shortest distance of this gene set within the interactome is 1.63. (A list of genes, as well as GEXF and results files for this gene set, are available in the supplementary materials.)

**Figure 1:**
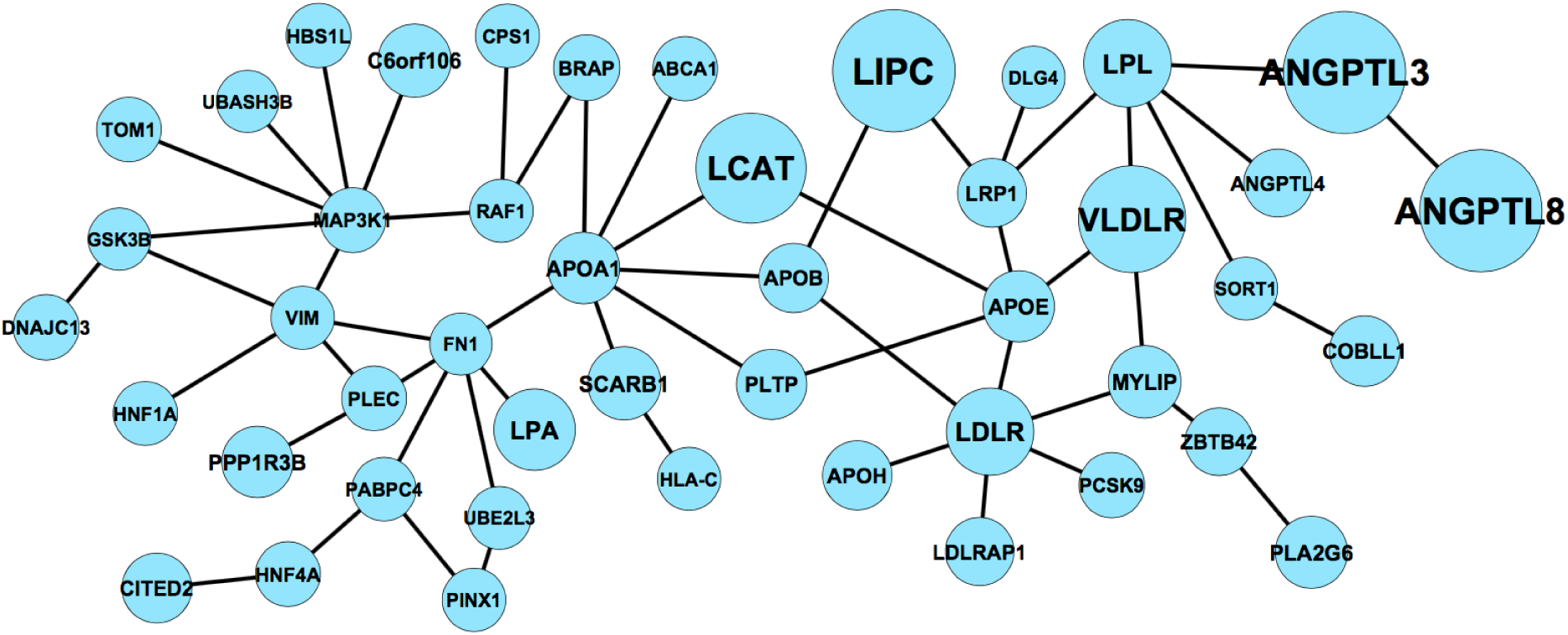
The largest connected component (LCC) of the gene set in the interactome. Each node is sized proportionally to its weight, which is the ratio of its degree in the gene set to its degree in the interactome.

## 4 Discussion

Inter-Tools is a simple-to-use command-line Python toolkit for assembling an interactome dataset and graph files of gene networks. It lowers the barrier-to-entry of interactome analysis by giving researchers easy access to an up-to-date interactome suited to their particular research needs, as well as a basic visualization tool. Of course, scientifially valid interactome analysis cannot be done exclusively visually; conclusive interactome analysis must be specifically designed for the question that the research is trying to answer. However, Inter-Tools provides a place to begin. It is a resource for interactome-based hypothesis generation and preliminary investigation.

